# Exploring the Winner-Loser Effect on Emotion in Crickets

**DOI:** 10.1101/2025.10.24.684431

**Authors:** Mason Miller, Elena Liebl, Michael S. Reichert

## Abstract

Individual variation in behavioral traits, commonly referred to as ‘personality,’ is exhibited in numerous animal species, including invertebrates. The specific personality trait of exploration tendency may be affected by positive/optimistic or negative/pessimistic emotional states. Animal contests can induce these emotions, as winning against one’s opponent increases ‘optimism’, while losing causes ‘pessimism’; these states can then affect behavior in other contexts. Previous studies demonstrate that house crickets (*Acheta domesticus*) experience this winner-loser effect in fights, thus we investigated if a cricket’s exploratory behavior is influenced by the outcome of its contest. We predicted that after a fight, winners would exhibit increased exploratory behavior due to an optimistic emotional state, while losers would show decreased exploratory behavior due to a pessimistic emotional state. Exploration tendency was measured as an individual’s latency to exit a shelter. Crickets then participated in an aggression contest, with opponents chosen to be smaller or larger so that focal individuals either won (n=33) or lost (n=31), respectively. Latency to exit was again measured after the contest, as well as 48 h later to examine the persistence of winner-loser effects on exploration behavior over extended time periods. Latencies for control crickets (n=40) were measured at the same time points, but controls did not engage in a fight. In one of two years, winning crickets’ latency times decreased following the contest, while losing crickets’ latency times were not significantly affected. This suggests that individuals’ emotional states were changed by the experience of winning a fight, which affected exploration tendencies, while losing a fight does not have an effect on exploration tendency. These data on invertebrate emotions have important implications for human health by allowing for future studies on the mechanisms behind mental states.

## INTRODUCTION

An animal’s experiences throughout its lifetime influence its behavior (Carlson, 2017), and individual variation in behavior can arise due to variation in internal states (Hertel et al., 2020). Two major components of internal states are valence and arousal (Stanley & Meyer, 2009). Valence states are characterized as positive or negative internal states induced by stimuli, and arousal is the intensity of the internal state (Mendl et al., 2010). These internal states can cause an individual to make judgments about external stimuli, resulting in changes in behavior and allowing the valence state to be observed (Crump et al., 2020). The judgment bias test (or “cognitive bias test”) has been utilized in previous research to assess the internal states of invertebrates (Solvi et al., 2016; Bateson et al., 2011). Individuals are trained to discriminate between distinct cues: one associated with a positive outcome (reward) and one associated with a negative outcome (punishment). Then, individuals are exposed to an intermediate or ambiguous cue. An individual with a positive valence state may indicate more ‘optimistic’ assumptions about the ambiguous stimuli if the response is similar to the positive or rewarding cue. On the other hand, an individual with a negative valence state may indicate more ‘pessimistic’ assumptions about the ambiguous stimuli by responding similarly when presented with the negative or punishing cue (Perry & Baciadonna, 2017; Mendl & Paul, 2020). These internal, short-term states induced by stimuli can be defined as emotions (Anderson and Adolphs, 2014; Perry & Baciadonna, 2017) and can be relevant to any animal with a nervous system. Emotional states have been investigated in another study without utilizing the cognitive bias test by examining individual judgment of novel stimuli and differences in valence in rodents (Verjat et al., 2020). After being exposed to positive or negative cues, individuals made decisions about novel stimuli that indicated an ‘optimistic’ or ‘pessimistic’ internal state based on willingness to approach and investigate. However, this individual judgment bias has not been well-studied in invertebrates to determine if internal states may affect behavior in other contexts.

An experience that has been shown to affect behavior in invertebrates is winning or losing an aggressive contest (Rillich & Stevenson, 2011). The outcome of prior contests can affect an individual’s future performance, such that winners have an increased chance of winning in the future, while losers have an increased chance of losing (Kasumovic et al., 2010). This phenomenon is referred to as the winner-loser effect, and the mechanism behind it is not well understood (Hsu et al., 2005). One possible explanation is that the mental states of individuals are altered by winning or losing, which causes a change in valence state and subsequent behaviors (Crump et al., 2020). By winning, an individual has a positive or ‘optimistic’ internal state about potential contests, similar to experiencing a reward. By losing, an individual has a negative or ‘pessimistic’ internal state about potential contests, similar to experiencing a punishment. If the impacts of the winner-loser effect on future contest behavior are due to emotional states, then contest experience could carry over to affect behavior in other contexts. To determine this, we examined the winner-loser effect in the house cricket, *Acheta domesticus*, as this species’ contest behavior has been well-studied and shown to exhibit the winner-loser effect (Hack, 1997). We predicted that the valence effects of a contest experience will be demonstrated by changes in behavior in another context, specifically the personality trait of exploration tendency.

Personality, or consistent among-individual differences in behavioral traits, has been increasingly studied in animal species (Dall, Houston, & McNamara, 2004), yet few studies have investigated personality in invertebrates (Kralj-Fišer & Schuett, 2014). These traits are characterized by behavior in one context correlating with the same or another behavior in a different context (Wolf & Weissing, 2012), such as aggressive individuals tending to be bolder in response to predators (Sih et al., 2004). One behavioral trait that has frequently been identified as a personality trait is exploration tendency, or the propensity to move about an unfamiliar environment (Hughes, 1997; Réale et al., 2007). Exploration tendency has been shown to be repeatable (Albers & Reichert, 2022), but is also a plastic trait that can be affected by experience or environment (Reader, 2015; Thompson et al., 2018). Thus, the exploration tendency of an individual may be consistent until it is affected by an experience, such as participating in a contest. A change in internal state or valence could be observed through measurable differences in an individual’s exploration tendency before and after a contest. The costs of exploration include time, energy, and predatory risks, but individuals benefit from obtaining information and resources if they choose to explore (Reader, 2015). If an individual won a contest and had an ‘optimistic’ internal state, it may be more exploratory due to interpreting an unfamiliar environment as an opportunity to gain information and resources. On the other hand, a losing individual with a ‘pessimistic’ internal state may interpret an unfamiliar environment as costly and explore less. This would allow for evidence regarding the potential for valence states caused by the winner-loser effect.

In this study, we tested the effects of winning or losing a contest on the exploration tendency of *A. domesticus*. We first performed a repeatability analysis of exploration behavior to determine if it is a personality trait, as suggested by previous studies (Albers & Reichert 2022). We then tested changes in valence state by examining differences in exploration tendency between winning and losing individuals before and after a contest. We predicted that individuals that win in a contest will have a positive or ‘optimistic’ valence state and an increase in exploration tendency, while individuals that lose in a contest will have a negative or ‘pessimistic’ valence state and a decrease in exploration tendency.

## METHODS

### Subjects

We used mature male *A. domesticus* individuals that were commercially purchased as 6-week-old juveniles from Fluker Farms. Individuals were kept in a group container that was 65.4 × 46.7 × 33.7 cm with a lid made partly of mesh material for oxygen flow. The container housed approximately 500 crickets. Fluorescent lighting was placed directly above the group container on a 12:12 h light:dark cycle. Egg cartons were utilized as shelters in the group container, and crickets were provided with water vials and fed chicken feed ad libitum (NatureWise Layer Feed, 16% Protein Pellet). Food and water were changed twice per week. 72 h prior to the first trial, sexually mature male crickets were separated and individually placed in transparent plastic containers measuring 20.32 × 20.32 × 7.63 cm. The individual containers each had a piece of egg carton, a water vial, and a petri dish of food. The crickets were isolated to ensure that there were not any prior aggressive contests in the group containers influencing the individuals in the trials (Adamo & Hoy, 1995). Male crickets were exclusively used in the trials due to their increased likelihood to engage in aggressive contests when exposed to a conspecific (Hack 1997).

### Experimental Design

A timeline of the experimental design is shown in **Figure 1**. Focal crickets were measured for initial latency to emerge from a shelter into a novel arena and randomly assigned to treatment groups (see *Aggression Trials* below for details). 24 h later, focal individuals participated in an aggression contest, with larger crickets expected to be winners and smaller crickets expected to be losers (Hack 1997). Following the contest, focal crickets were immediately measured for post-treatment latency to emerge. We then measured latency to emerge 48 h after the contest to test whether contest experience has long lasting effects on behavior. The latency to emerge trials were conducted and measured in the same way for control individuals, but they did not compete in a contest.

**Figure 1.**
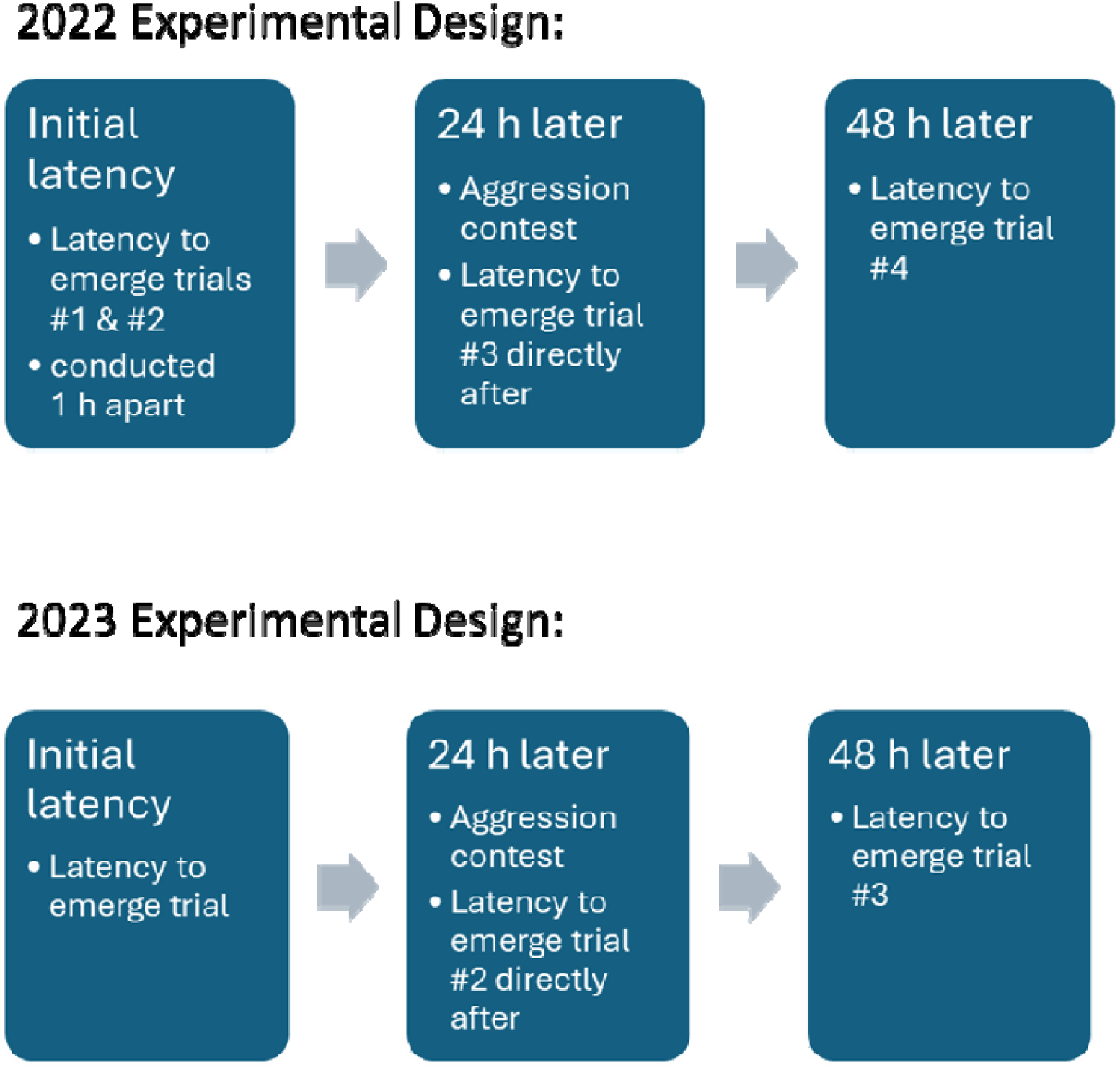
Timeline for experimental design. Trials in 2022 differed in the number of latency to emerge trials compared to 2023 due to testing repeatability.

In 2022, our sample included winners (n=20), losers (n=17), and controls (n=20). After reviewing the data, the control treatment individuals unexpectedly displayed significantly lower latency to emerge measurements when compared to the other treatment groups. This may be in part due to the trials for the control group being conducted at a later date after all winner and loser focals were tested. We therefore conducted further trials in 2023 with winners (n=13), losers (n=14), and controls (n=20). The experimental design was identical in the two years except that in 2022 we performed four latency to emerge trials: two trials conducted 1 h apart before the contest to test for repeatability, one trial directly following the aggression contest, and one trial 48 hours later. However in 2023 we performed only three latency to emerge trials: one trial 24 h before the aggression contest, one trial directly after the contest, and one trial 48 h post-contest. We only used one pre-contest trial in 2023 because we had already established in 2022 that latency to emerge is repeatable. In 2023, the control individuals were tested along with the winners and losers instead of at a later date to try to mitigate any confounding effects.

### Latency to Emerge

Individuals were placed at the bottom of an opaque, plastic cup (5 cm diameter, 7.6 cm height), which served as the shelter. The shelter was positioned in the same corner of an empty opaque container measuring 37.4 x 25.4 x 13.4 cm. The trial began once the cup was placed horizontally in the container with the opening facing outwards into the arena. We recorded latency to emerge as the time in seconds that it took for the cricket to place all legs outside the shelter, indicating it had left the shelter. Each individual was given up to 600 s to emerge from the shelter. If the focal cricket did not leave within 600 s, the trial was terminated and its latency score was recorded as 600 s. The arena was cleaned with ethanol between each trial. Both focal and control individuals participated in the latency to emerge trials.

### Aggression Trials

Sexually mature male crickets were designated as focals or opponents. Focal individuals participated in the latency to emerge trials and had their aggression scores calculated. Opponent individuals did not participate in latency to emerge trials and were only utilized in the aggression contests to provide an opponent for the focal individual. Focal individuals were randomly paired with an opponent that differed by approximately 0.1 grams in body weight. The average body weight for focal individuals was 0.27 grams in 2022 and 0.2 grams in 2023. Control individuals had a similar body weight as the focal crickets and were handled in the same way but did not participate in an aggressive contest. If the focal cricket was assigned to the loser treatment, then it was paired against a heavier opponent. If the focal cricket was assigned to the winner treatment, then it was paired against a lighter opponent. Each focal cricket only competed in one aggression contest. In 2022, each opponent cricket competed 2-3 times, facing multiple focal individuals. The opponent was given at least 10 min between each contest to allow them to recover aggressive motivation. However, in 2023, each opponent was only utilized in one contest.

Aggression trials consisted of each focal and opponent cricket being placed in a circular, plastic arena (10.8 cm diameter, 7.6 cm height) at the same time. To acclimate to the arena before the contest began, each cricket was placed underneath a separate plastic cup in the arena for 2 min. At the end of the acclimation period, the cups were removed and the crickets were allowed to interact for 5 min. The focal cricket was designated by a small mark of red paint on the abdomen to allow for recognition during video playback and contest scoring. Their interactions were recorded with a Panasonic HC-V770 Camcorder. The outcome of each contest was also noted. Following each aggression contest, the arena was cleaned with ethanol after the crickets were removed. The aggression scores for the focal individuals were quantified through the observation of nine different behaviors, based on Adamo & Hoy (1995) and Albers & Reichert (2022).

### Statistical Analyses

#### Repeatability of exploration tendency pre-contest

We estimated the repeatability coefficients for latency to emerge using the ‘rptR’ v.0.9.22 package (Stoffel et al., 2017) in R v.3.6.1 software (R Core Team, 2021). In the 2022 trials, repeatability for latency to emerge was estimated with the time to exit the shelter as the (Gaussian-) dependent variable, first and second trial numbers as a fixed effect and cricket identification (ID) as a random effect (N = 57).

#### Effects of contest experience on latency to emerge

To test the relationship between trial number and latency to emerge in both the 2022 and 2023 trials, we created separate models for each year using the ‘lmer’ function in the ‘lme4’ v.1.1-28 package in R as a linear mixed model with latency to emerge as the (Gaussian-) dependent variable, and trial number and interaction between trial number and treatment as fixed effects. Cricket identification was a random effect. For the short-term effects of contest experience on latency to emerge, we looked at the difference in latency to emerge between the trial before the contest and the trial directly following the contest. We used separate models for winners only, losers only, and both. To assess the long-term effects of contest experience on latency to emerge, we also included a fixed quadratic effect of trial number.

## RESULTS

### Repeatability of Latency to Emerge Behavior

In 2022, the latency to emerge from the shelter was significantly repeatable (R=0.45, CI [0.24, 0.63], p=0.000145) (**Figure 2**).

**Figure 2.**
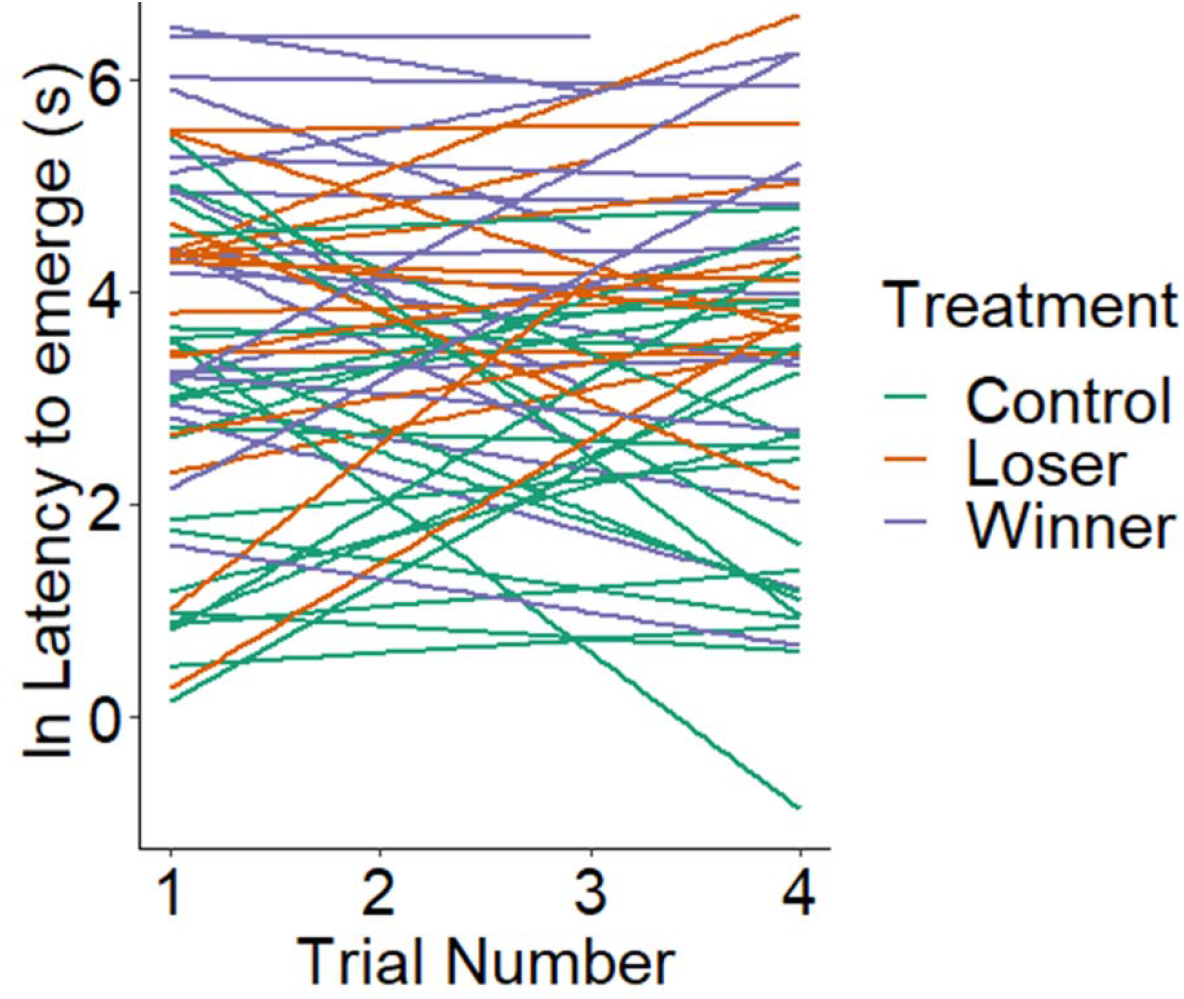
Repeatability of latency to emerge. Each line represents an individual focal or control cricket from the 2022 latency to emerge trials.

### 2022 Effects of Contest Experience on Latency to Emerge

There was a significant interaction between trial number and treatment (t=-2.931, p=0.006) for the analysis across all trials (latency to emerge before the contest, after the contest, and 48 h after). There was also a significant interaction between trial number and treatment for the analysis that only included the trial directly before the contest and directly after the contest (t=-2.899, p=0.00643). Focal individuals designated as winners (n=20) decreased significantly in latency time (analysis for winner treatment only; t=-3.295, p=0.00381) after participating in an aggression contest, while focal individuals designated as losers (n=17) did not significantly change their latency time (analysis for loser treatment only; t=0.905, p=0.3791) after participating in the contest (**Figure 3**). Control individuals (n=20) did not exhibit a difference in latency time (analysis for control treatment only; t=0.06, p=0.95) across all trials. There was a significant interaction between squared trial number and treatment for latency to emerge (t=2.892, p=0.00524): winners returned to pre-fight behavior in trial 4.

**Figure 3.**
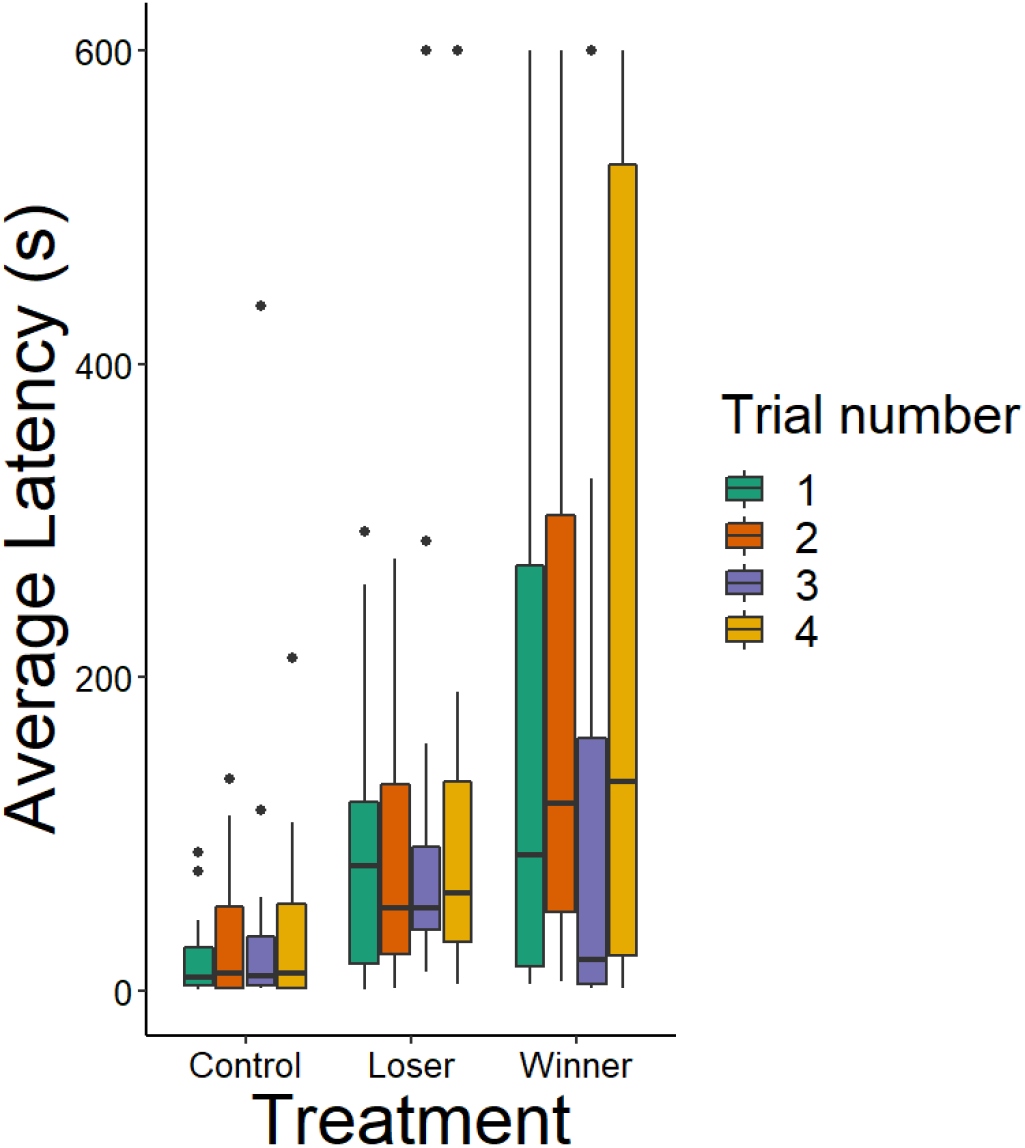
Relationship between latency to emerge, trial number and treatment for all trials in 2022. Contests occurred between trials 2 and 3. Outliers are characterized as points.

### 2023 Effects of Contest Experience on Latency to Emerge

There was not a significant interaction between trial number and treatment (t=0.019, p=0.99) for the analysis across all trials. There was also not a significant interaction between trial number and treatment for the analysis that included only the trial directly before the contest and directly after the contest (t=-0.645, p=0.525). Focal individuals designated as winners (n=13) appear to have increased in latency following an aggression contest **(Figure 3)**, but not significantly so (analysis for winner treatment only; t=-0.656, p=0.52408). Losers (n=14) appear to have a decrease in latency time (**Figure 4**) but this decrease was also not significant (analysis for loser treatment only; t=0.113, p=0.912). Control individuals (n=20) did not exhibit a significant difference in latency time (analysis for control treatment only; t=-1.59, p=0.12) across trials. There was not a significant interaction between squared trial number and treatment for all trials (t=0.399, p=0.6919).

**Figure 4.**
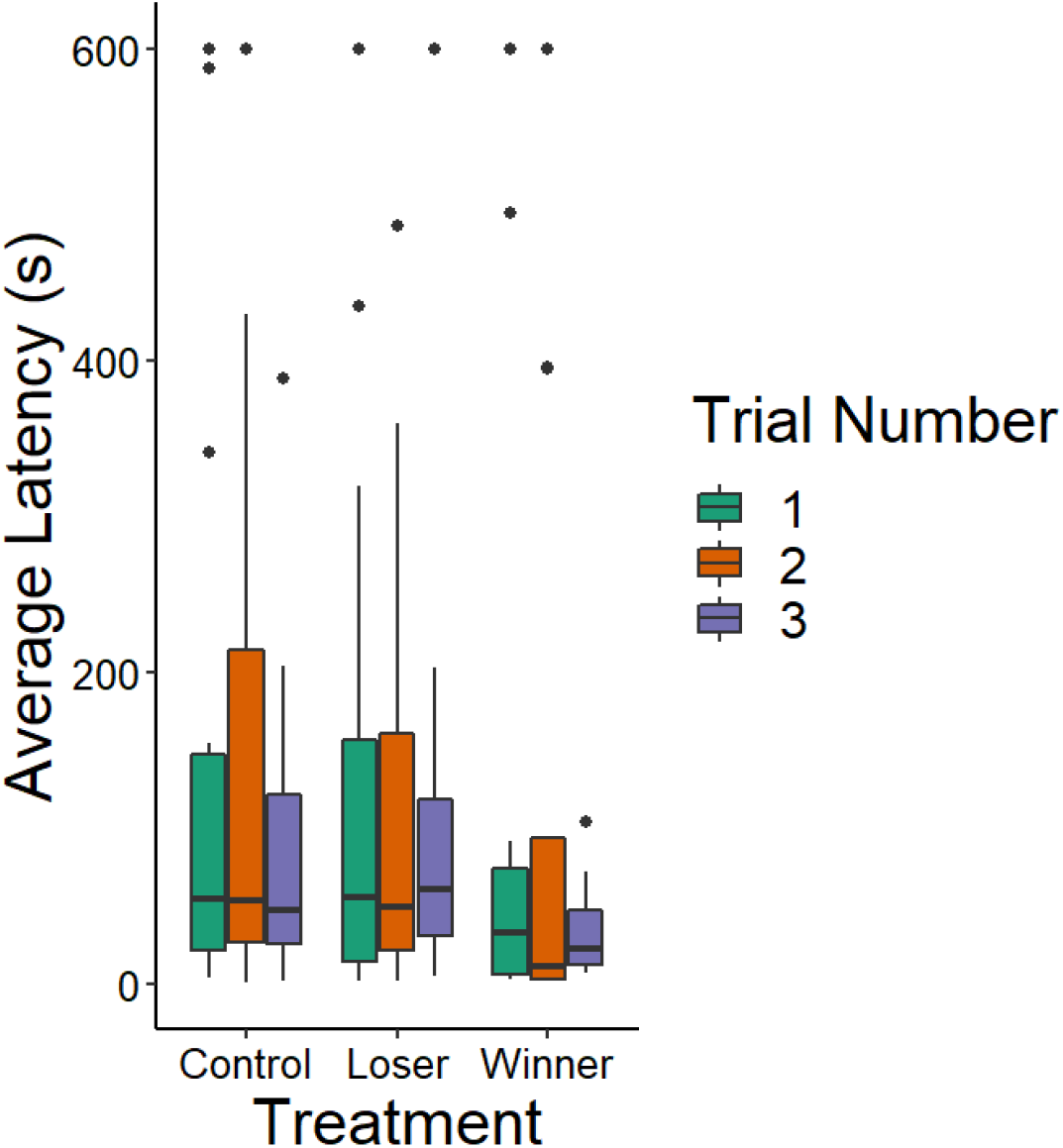
Relationship between latency to emerge, trial number and treatment for all trials in 2023. Contests occurred between trials 1 and 2. Outliers are characterized as points.

## DISCUSSION

As expected, *Acheta domesticus* exhibited decreased latency to emerge from the shelter after winning a contest in the 2022 trials. Individuals that won took less time to exit the shelter into the arena, displaying more exploratory behavior. However, focal crickets that were winners in the 2023 trials did not show a significant difference in latency to emerge before and after the contest. On the other hand, individuals that lost a contest did not show significantly increased latency to emerge following a contest in either the 2022 or 2023 trials. These data did not align with our hypothesis as we predicted that losing a contest would impact latency to emerge and thus, exploration tendency. Individuals’ exploration tendency before the contest was repeatable as shown in the 2022 trials, which corroborates the findings of Albers & Reichert (2022). The control individuals had consistent exploration tendency across the trials, suggesting that the contest was the experience causing differences in pre-and post-contest latency to emerge. 48 h after the contest, the winning crickets’ latency to emerge increased in both 2022 and 2023, indicating a return to the pre-contest internal state.

Based on these results, experience winning a contest has an effect on exploration tendency, while losing a contest does not have a significant effect. In either case, future research is necessary to support these findings as results from 2022 and 2023 differed for the winners. Furthermore, the visualized data for the losing individuals indicates that latency to emerge may have decreased in the 2023 trials similar to the winners, which would need further research to investigate due to a lack of significant findings from our study. A possible explanation for the decrease in latency to emerge from the losers could be that individuals with a ‘pessimistic’ internal state attempt to alleviate the uncertainty of their surroundings by exploring more (Verjat et al., 2020).

There were several changes in the methods between the 2022 and 2023 trials that may have caused differences in the findings. In 2022, focal individuals competed against opponents that had been utilized in multiple contests. While the opponent individuals were allowed a rest period, there may have still been winner-loser effects carried over to the next contest they participated in. This could have caused the opponents to have a higher state of arousal or more intense aggressive behaviors if they were winners, or a decreased state of arousal or less intense aggressive behaviors if they were losers. The intensity of the aggression of the opponents in 2022 may have caused a different outcome of the contest compared to the 2023 when each opponent was paired with a focal and only utilized in one contest. Another difference was the timeline of the testing of the control individuals. In 2022, controls were tested after all of the focals were tested, while in 2023, controls were tested each week along with the focals. Due to this variance, possible confounding variables may have been differences in the time of day of testing, age of the crickets, or environment.

More evidence regarding correlations between contest behavior and behavior in other contexts would allow for a better understanding of the mechanism underlying the winner-loser effect. Future directions of this study could include investigating the effects of winning or losing an aggression contest on other personality-dependent behaviors, such as sociability (Kaiser & Müller, 2021). Furthermore, emotional states as quantifiable changes in behavior may be engaging specific neurobehavioral systems that are linked to experiences (Mendl et al., 2010), but more research is needed to understand these systems. *A. domesticus* could be an ideal model specimen for future studies regarding the mechanism behind emotions as crickets’ simple neural architecture provides opportunities for examining the specific pathways of changes in emotional states (Perry & Baciadonna, 2017). Furthermore, in other invertebrates such as honeybees, the activity of a single neuron can be associated with a specific behavior like reward learning (Hammer, 1993). Investigating if cricket neural pathways correlate with specific behaviors in the context of ‘optimism’ and ‘pessimism’ could allow for the examination of the mechanisms underlying changes in emotional states. Gathering more data on valence could yield results relevant to the understanding of human mental health. In humans, pessimistic individuals are more likely to experience depression and more severe episodes (Schueller & Seligman, 2008).

Thus, data on the underlying mechanisms of pessimism and optimism in invertebrates may be useful for future research on the helpful manipulation of human mental states through advanced therapeutic techniques and treatments.

## ACKNOWLEDGEMENTS

We thank Kennedy Funa and Kayleen Sugianto for help with the experiments and members of the Reichert lab for feedback. Funding was provided by a grant from the Lew Wentz Foundation to M.M.

